# Long-Term Imaging of Cellular Forces with High Precision by Elastic Resonator Interference Stress Microscopy

**DOI:** 10.1101/118554

**Authors:** Nils M. Kronenberg, Philipp Liehm, Anja Steude, Johanna A. Knipper, Jessica G. Borger, Giuliano Scarcelli, Kristian Franze, Simon J. Powis, Malte C. Gather

**Affiliations:** SUPA, School of Physics and Astronomy, University of St Andrews, St Andrews, UK; Institute of Immunology and Infection Research, University of Edinburgh, Edinburgh, UK; Fischell Department of Bioengineering, University of Maryland, College Park, MD, USA; Department of Physiology, Development and Neuroscience, University of Cambridge, Cambridge, UK; School of Medicine, University of St Andrews, St Andrews, UK

## Abstract

Cellular forces are crucial for many biological processes but current methods to image them have limitations with respect to online analysis, resolution and throughput. Here, we present a robust approach to measure mechanical cell-substrate interactions in diverse biological systems by interferometrically detecting deformations of an elastic micro-cavity. Elastic Resonator Interference Stress Microscopy (ERISM) yields stress maps with exceptional precision and large dynamic range (2 nm displacement resolution over a >1 μm range, translating into 1 pN force sensitivity). This enables investigation of minute vertical stresses (<1 Pa) involved in podosome protrusion, protein specific cell-substrate interaction and amoeboid migration through spatial confinement in real time. ERISM requires no zero-force reference and avoids phototoxic effects, which facilitates force monitoring over multiple days and at high frame rates and eliminates the need to detach cells after measurements. This allows observation of slow processes like differentiation and further investigation of cells, e.g. by immunostaining.

## Introduction

Cells sense mechanical cues and constantly probe the mechanical properties of their environment (mechano-sensing)^1^. Mechanical stimuli at the cellular level play an important role in cell function, e.g. in migration and differentiation, as well as in tissue-level processes, e.g. morphogenesis, tumour growth and immune response^2–7^. Forces exerted by cells are fundamental not only for mechano-sensing but also for many other physiological processes such as locomotion (e.g. during immune response or tumour metastasis), cell growth, wound healing, tissue formation and repair, and extracellular matrix deposition^8–14^.

The mechanical interaction between a cell and its substrate can occur through various mechanisms. During mesenchymal migration, lateral mechanical forces generated by the cytoskeleton are transferred to the extracellular matrix via firm focal adhesion protein complexes^15^. In other cases forces act perpendicularly to the substrate (vertical forces) and there is now a rapidly increasing interest in the role vertical forces play, e.g. during the extension of special cellular processes such as podosomes^16^ or during amoeboid migration in spatial confinement where non-specific cell-substrate interactions such as friction and pushing are important^17–19^. The forces involved in these diverse mechanical interactions are expected to vary substantially in magnitude, spatial distribution and temporal evolution. Being able to monitor them in a robust and non-disruptive manner is thus critical to advancing our understanding of these processes.

Early studies in this field obtained semi-quantitative maps of the applied forces by direct observation of wrinkles that cell generated forces induce in elastomer substrates^20,21^. Currently, the most widely used methods for imaging cellular forces (i.e. to record stress or force maps) are traction force microscopy (TFM)^22–28^ and the use of micro-fabricated elastic micro-pillar arrays^8,29,30^. (Ref. 31 reviews some of the presently available tools for measuring cell generated forces.) TFM and micro-pillar arrays are based on tracking the displacement of fluorescent particles in hydrogels and the bending of individual elastic pillars, respectively. While being powerful, these methods are indirect; the data in most cases need to be analysed offline, and if fluorescence is used, this can be associated with photo-toxicity. Furthermore, zero-force reference images are usually needed if cell-induced substrate deformations are smaller than approximately 200 nm^29,32^. For these reference images cells have to be removed from the substrate, rendering them unavailable for further investigations. There have been attempts to obviate the need for disruptive cell removal in TFM by using highly regular meshes or grids to record cell induced substrate deformations but even under optimized conditions these approaches are limited to detecting relatively large deformations and require fluorescence microscopy as well as rather involved sample preparation^32–34^. Micro-pillar arrays and TFM also have limited sensitivity if the forces of interest are exerted perpendicularly to the substrate. For instance, measuring the weak vertically-directed protrusive forces exerted by podosomes has necessitated a specialized detection scheme based on atomic force microscopy (AFM)^16^. What is currently missing is a more generally applicable method to image cellular forces that allows cells to be retained on the substrate for subsequent measurements and is capable of resolving weak forces with reasonable throughput.

Here, we address this need by introducing Elastic Resonator Interference Stress Microscopy (ERISM), a new approach for online, robust and non-destructive imaging of forces associated with various types of mechanical cell-substrate interactions. While most existing methods use localization microscopy or direct imaging of surface deformations, ERISM is based on interferometrically detecting cell-induced substrate deformations by using an elastic optical micro-resonator and thus provides unprecedented sensitivity. We are able to resolve not only forces exerted by cells that form firm, integrin-based, focal adhesion contacts to the substrate but can also detect protein-specific cell-substrate interaction and quantify the much weaker vertical forces (down to piconewtons) that are associated with amoeboid-type cell migration through confined environments and with the protrusion of podosomes. ERISM requires no zero-force reference image, which eliminates the need to detach non-migrating cells after a measurement and enables continuous, long-term measurements of multiple cells on one substrate as well as further investigation of the cells, e.g. by immunostaining. Being a wide-field imaging method, ERISM determines the local deformation at each point of the image simultaneously and requires only low light intensities, thus facilitating observation of multiple cells at once without inducing photo-damage to the cells.

## Results

### Concept and validation of ERISM

Cells were grown on a protein-coated elastic optical micro-cavity consisting of a layer of an ultra-soft siloxane-based elastomer (stiffness 1.3 kPa, thickness (8±0.5) μm; unless stated otherwise) sandwiched between two thin and semi-transparent gold layers (**Fig. 1a, b**; see **Supplementary Fig. 1** for details on the gold-elastomer interface). Forces applied by cells were mapped with the microscope setup schematically shown in **Figure 1c.** Cells of interest were identified by conventional bright-field, phase contrast or fluorescence microscopy (**Fig. 1d**). The reflection of the micro-cavity was then imaged under illumination with low-intensity monochromatic light (**Fig. 1e, Supplementary Fig. 2**). When a cell locally deformed the micro-cavity, the incident light coupled to resonant modes of the micro-cavity at positions within the field of view where the micro-cavity thicknesses fulfilled a resonance condition and the reflectance at these positions was hence reduced (dark fringes). This fringe pattern provided a direct measure of how much the cells deformed the substrate, with each fringe corresponding to a deformation of about 200 nm. To obtain a more accurate map of the local deformation, reflectance images were recorded for a set of different illumination wavelengths. This yielded the local reflectance as a function of wavelength (**Fig. 1f**) and thus the local resonance wavelengths of the micro-cavity for every pixel within the field of view. The actual thickness for each pixel was then obtained by fitting the measured resonance wavelengths with an optical model using a fast look-up table-based fitting algorithm (**Supplementary Fig. 3**). A background plane was subtracted from the measured thickness map to obtain the cell-induced deformation (**Fig. 1g**).

**Figure 1.**
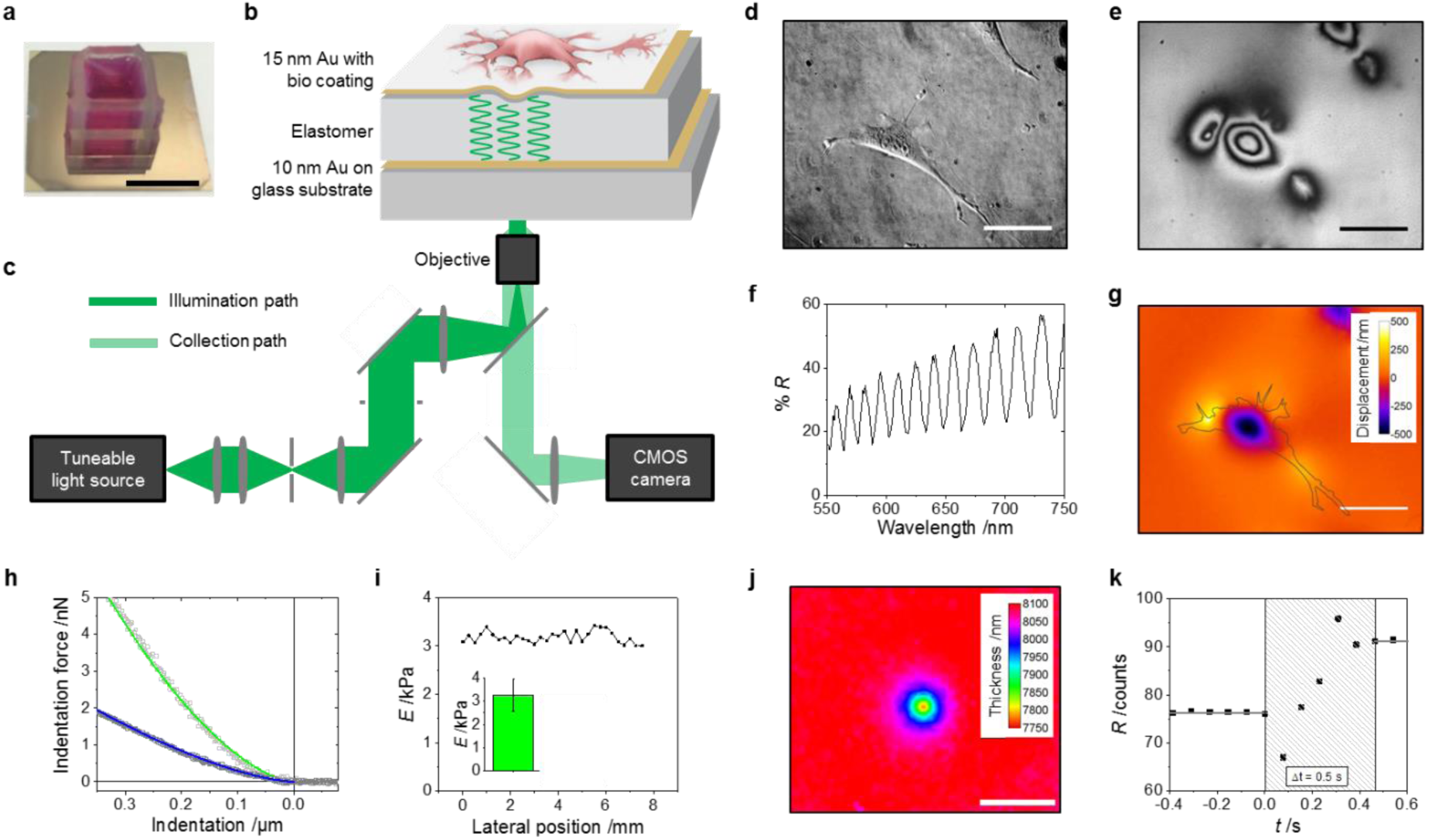
Elastic Resonator Interference Stress Microscopy (ERISM). (a) Photograph of the micro-cavity with a silicone chamber filled with cell medium mounted on top. (b) Sketch of layered structure of the micro-cavity with elastomer layer sandwiched between two ultra-thin gold mirrors. (c) Schematic of the wide-field optical readout system with tuneable light source providing monochromatic illumination and epi-collection to record spatially resolved images of micro-cavity reflectance. (d) Phase contrast image of a cell attached to the micro-cavity surface. (e) Co-registered monochromatic reflectance image of micro-cavity at 655 nm wavelength of illumination. (f) Wavelength dependence of the reflectance for one representative pixel within the field of view. (g) ERISM map obtained from the reflectance data, showing deformation induced by cell from (d). Outline of cell in black. (h-k) Validation of ERISM by atomic force microscopy (AFM). (h) Force-distance curves (open symbols) obtained by AFM micro-indentation with a 17 μm-diameter glass sphere and corrected Hertz model fits to the data for pristine elastomer (blue line) and for complete micro-cavity (green line). (i) Apparent stiffness obtained from Hertz model fit along the surface of a micro-cavity. Inset: Mean stiffness and standard deviation for micro-cavities produced in different batches (five independent measurements). (j) ERISM thickness map of micro-cavity upon AFM micro-indentation with a 17 μm-diameter glass sphere (set-force, 5 nN). (k) Reflectance change of elastic micro-cavity when quickly (10 ms) removing the AFM indenter from a 600 nm indentation. Scale bars, (a) 1 cm, (d, e, g) 50 μm, (j) 25 μm.

We used AFM to characterize the mechanical properties of the elastic micro-cavities. The apparent stiffness of the micro-cavities was determined by fitting a corrected Hertz model to force-distance data from micro-indentation measurements (**Fig. 1h**)^35^. Micro-cavities used in this study had an apparent stiffness of 3.2 kPa, in the same range as many relevant types of tissue^36^. The stiffness variation across the surface of a cavity was 0.12 kPa (standard deviation of 31 measurements across cavity surface, **Fig. 1i**), and the variation between cavities fabricated in different batches was 0.69 kPa (standard deviation for 5 batches, inset **Fig. 1i**). **Figure 1j** shows an ERISM thickness map for a micro-cavity into which an AFM probe was indented with a set-force of 5 nN. Using the stiffness measured by AFM, finite-element-method (FEM) modelling was used to convert this indentation profile – or later, cell induced deformations – into a map of locally applied stress (see Online Methods). Integration over the entire area of the indentation provided an estimate of the total applied force; for the AFM indentation above, the FEM model estimated an applied force of 4.3 nN, i.e. within 20% of the nominally applied force. By analysing the width of the indentation in **Figure 1j**, we estimated that the *lateral* resolution of ERISM is 1.6 μm (**Supplementary Fig. 4**), which is similar or better than values achieved with super-resolution TFM^28^ and the latest generation arrays of sub-micrometre sized elastic pillars^30^. However, due to the interference-based measurement principle, ERISM can measure the cell induced vertical substrate deformation with much higher resolution; in the present study, the displacement resolution was limited by the surface roughness of the micro-cavity (root mean square roughness 2 nm, **Supplementary Fig. 4**). The temporal resolution, i.e. the relaxation response time of the elastic micro-cavity, was < 0.5 s (**Fig. 1k**). We also tested the mechanical stability of the micro-cavity under repeated AFM indentations (**Supplementary Fig. 4 and Supplementary Video 1**) and under standard cell culture conditions (i.e., cell culture medium, 37 °C, 5vol% CO_2_ atmosphere). No significant change in stiffness was found after repeated indentations (Δ*E*/*E*_0_ < 10% after 60 cycles with 0.6 μm indentation depth) and no swelling was observed over 6 days of ERISM recording (the longest time course we have continuously monitored). Micro-cavities were kept in the incubator with cells growing on them for more than two weeks without any sign of degradation.

To demonstrate the broad applicability, long-term stability and high sensitivity of ERISM, we next discuss mechanical measurements for several cell types and for different forms of cell-substrate interaction.

### Localized vertical forces of podosome protrusion

The forces macrophages apply during podosome protrusion are perpendicular to their substrate and too small to be resolved with TFM. Podosomes are small (<1 μm), cylindrical, actin-based structures on the ventral side of the plasma membrane that are formed by many different cell types, e.g. invasive cancer cells and macrophages^16^. They play a critical role in cell migration and invasion into tissue, and it has recently been shown that they perform mechanosensing by protruding into their substrate^16^. Here we exploited the unprecedented vertical resolution of ERISM to resolve the protrusive forces of podosomes.

**Figure 2a** shows a phase contrast and an ERISM image of a primary human macrophage on a collagen-coated ERISM micro-cavity. The ERISM data was dominated by an unstructured broad pushing of the main cell body into the substrate and pulling at the cell periphery. However, there is also a more subtle substructure underneath the cell, which we hypothesized may be associated with podosomes protruding into the substrate. To visualize this structure more clearly, we removed the broad features from the data through spatial Fourier filtering of components with spatial frequencies smaller than ≈0.5 μm^-1^ (see **Fig. 2a** bottom panel and **Fig. 2b**). This revealed a large number of weak and tightly confined pushing sites (peak stress ≈10 Pa; diameter, 1-2 μm) which are surrounded by even weaker partial rings of upward pulling. Immunocytochemistry suggested that these features indeed corresponded to podosomes (**Fig. 2g**, see below). Although the deformations, stresses and integrated forces exerted by the podosomes were substantially smaller than what is usually observed for mechanical cell-substrate interactions mediated by firm focal adhesion contacts of e.g. fibroblasts^22,28^, and even though the podosome indentations were often less than 2 μm apart, they were clearly resolved by ERISM.

**Figure 2.**
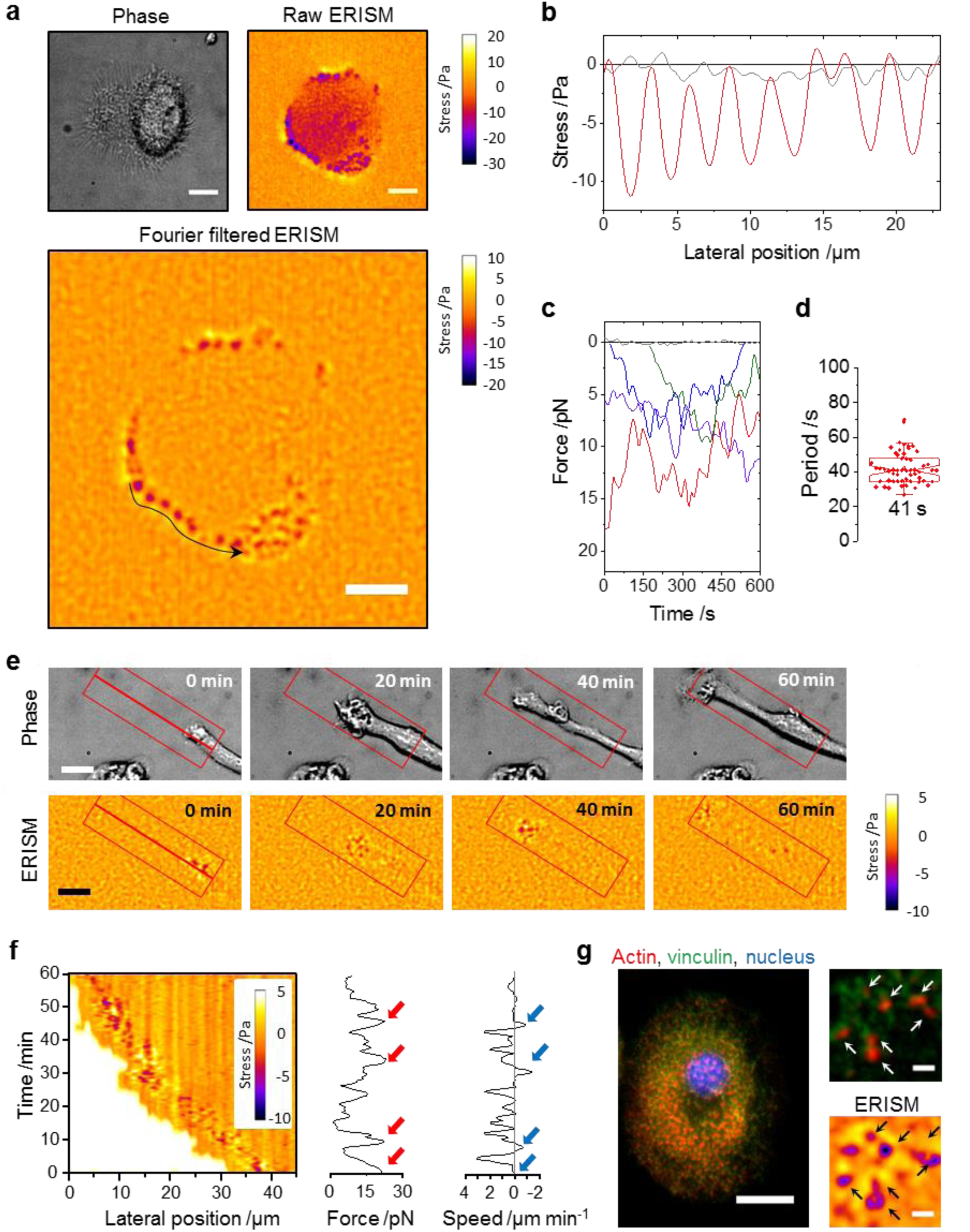
Investigation of podosome protrusions by ERISM. (a) Phase contrast (upper left), ERISM (upper right) and spatial Fourier-filtered ERISM image (bottom) of a macrophage. (b) Stress profile along the long black arrow in the Fourier filtered ERISM map in (a) (red line) and the typical noise in a region outside the cell area (grey line). (c) Temporal evolution of the force applied by several podosome protrusions and for a control area away from a podosome (grey line). Data extracted from a 1 h ERISM time-lapse measurement with a frame every 5 s. (d) Boxplot showing oscillation periods for 50 podosomes from five different cells. Each data point represents one podosome. (e) Investigation of podosome protrusion during cell expansion. Top row: Phase contrast images showing an elongating macrophage for four different time points. Bottom row: Corresponding Fourier filtered ERISM maps. (f) Left: Kymograph analysis of applied stress along bold red line in (e) averaged over a 7 μm-wide area above and below the line as illustrated by red rectangles in (e). White areas in kymograph are areas into which the cell has not yet expanded. Centre: Temporal evolution of the total force applied by all podosomes in leading edge of macrophage. Right: Temporal evolution of expansion speed of the cell. (g) Epi-fluorescence image of fixed macrophage with staining for actin, vinculin and nuclear DNA. Insets, magnified epi-fluorescence (top) and co-located ERISM map (bottom) with arrows indicating positions of individual podosomes. Scale bars in insets of (g) 2 μm, all other scale bars 10 μm.

Using time-lapse measurements, we also analysed the temporal evolution of the mechanical action of podosomes and found a pronounced oscillatory behaviour (**Supplementary Video 2**). **Figure 2c** shows the temporal evolution of the forces applied by several representative podosomes. (Force is calculated by integrating the applied stress over the area of the podosome.) We analysed a total of 50 podosomes from five different cells and found a characteristic period of the force oscillation of (41±10) s (**Fig. 2d**), consistent with earlier studies using a specialized AFM based detection scheme^16^.

We next investigated podosome behaviour during cell migration where their activity is believed to be most important^37^. **Figure 2e** shows phase contrast and Fourier filtered ERISM time-lapse images of the leading process of a macrophage (also see **Supplementary Video 3**). Over 1 h, the leading edge expanded by 40 μm. This was accompanied by the pronounced formation of protrusions in a region ≈5 μm away from the leading edge of the cell. **Figure 2f** shows a kymograph of the mean applied stress along the direction of cell migration (along the line shown in **Fig. 2e**). The forces generated by podosomes in the region around the leading edge of the cell oscillated over time (**Fig. 2f**; red arrows indicate times of maximum pushing). The oscillations were likely associated with the speed of cell expansion as maximum podosome activity was immediately followed by stalling of cell migration, and in some cases even by the retraction of the leading edge (negative speed, blue arrows).

Finally, to confirm that the observed structures were indeed podosomes, we fixed the cells on the micro-cavity immediately after recording an ERISM image and performed immunostaining for actin and vinculin. As ERISM can be readily combined with other imaging modalities, we then recorded the actin and vinculin distribution by epi-fluorescence imaging through the micro-cavity substrate. This revealed a characteristic podosome dot-pattern with small actin-rich spots surrounded by vinculin rich regions (the soft substrate led to some mislocalization of vinculin^16^). The actin-rich spots co-localized with pushing sites in the stress map and the surrounding vinculin-rich regions co-localized with the weaker ring-shaped pulling (**Fig. 2g**).

### Diffuse forces during amoeboid migration in confined environments and adhesion receptor specific cell-substrate interaction

While the podosome protrusions discussed above represent very localized cell-substrate interactions, other cell types, such as amoebae, transmit weak and more diffuse vertical stresses through transient, diffuse adhesion contacts when migrating in a three-dimensional environment ^17,38,39^. This “amoeboid migration” is also observed in embryonic cells, cancer cells and leukocytes^17,18,40^. It is characterized by rapid changes in cell shape, induced by the formation of pseudopodia and plasma membrane blebs. In spatially confined environments, friction or a “chimneying”-type of locomotion, in which cells project themselves forward in horizontal direction by applying weak vertical forces, have been proposed as mechanisms for stress transduction^17,18,38,40^. However, measurements of substrate displacements and the involved force patterns remain challenging, mainly due to the technical difficulties involved in detecting small vertical displacements with sufficient resolution.

We used ERISM to study deformations induced by migrating *Dictyostelium discoideum* amoebae confined within a 5 μm-thick void as schematically illustrated in **Figure 3a**. During migration, a continuous evolution of an ellipsoid push and pull pattern was observed (**Fig. 3b and Supplementary Video 4**). Profiles of the stresses along the longer axis of the cell are shown in **Figure 3c** to illustrate the horizontal cell migration more clearly. Importantly, horizontal migration occurs even though the detected peak stresses were <1 Pa.

**Figure 3.**
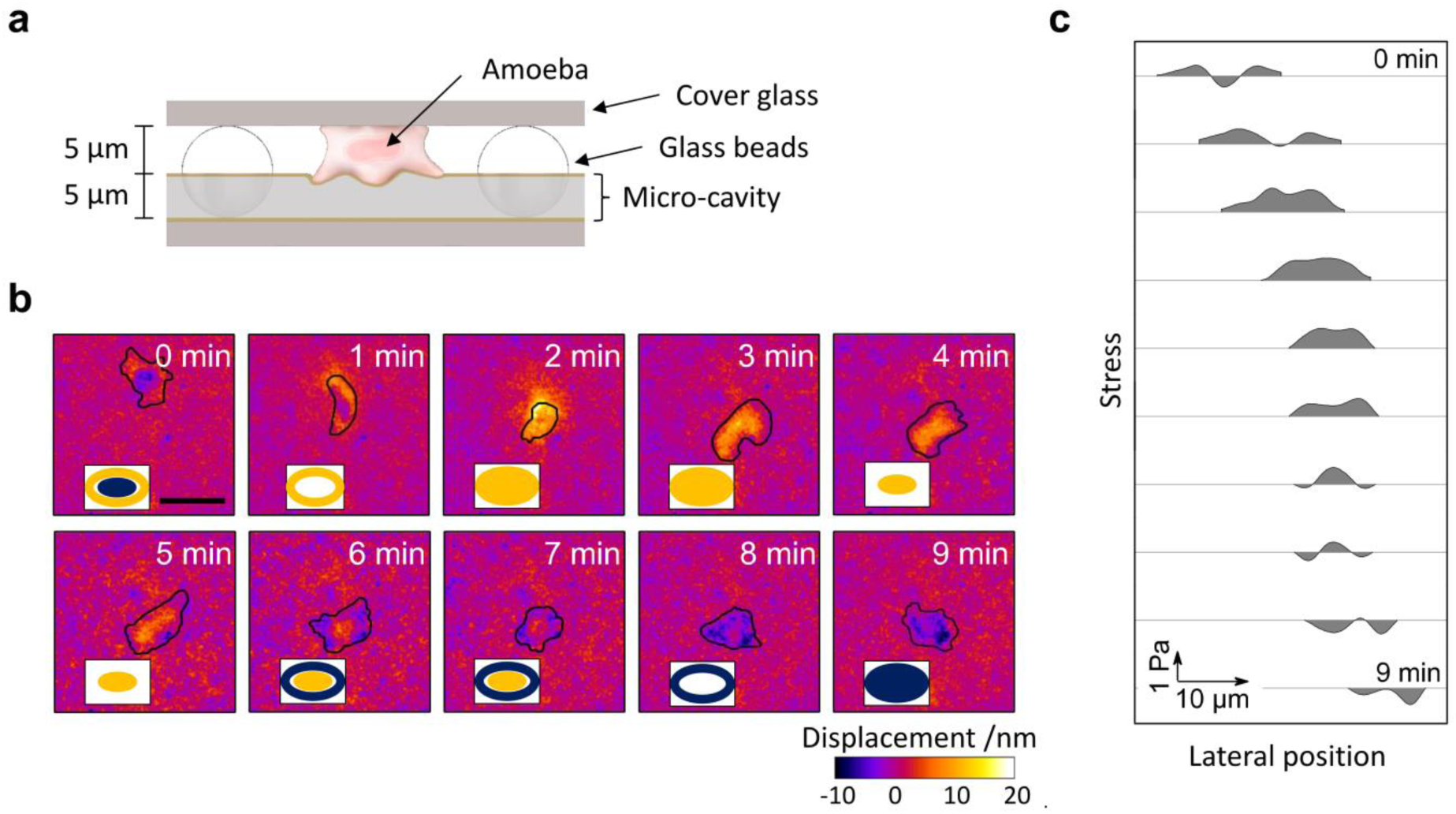
Investigation of amoeboid migration in confined space using ERISM. (a) Illustration of 5 μm void formed on top of a 5 μm thick micro-cavity substrate by using 10 μm-diameter glass beads and a glass coverslip. (b) Temporal evolution of substrate deformation generated by a *Dictyostelium discoideum* amoeba migrating through the void. The boundary of the cell is indicated in black. The insets show schematic illustrations of the displacement pattern at each time point. (c) Profiles of stress along the long axis of the cell (with 8 μm bandwidth spatial averaging to filter surface roughness). Scale bar 15 μm.

We also applied ERISM to perform wide-field imaging of adhesion receptor specific cell-substrate interaction, specifically during the adhesion of primary murine T cells. During lymphocyte migration and immunological synapse formation between T cells and antigen presenting cells, T cells adhere to the extracellular matrix and to other cells^42,43^. The adhesion is based on a protein complex formed between Lymphocyte function-associated antigen 1 (LFA-1) – a transmembrane integrin on the surface of T cells – and Intercellular Adhesion Molecule 1 (ICAM-1). We measured LFA-1:ICAM-1-mediated adhesion by investigating T cells on ICAM-1-coated micro-cavity substrates. During initial contact, a significant fraction of T cells deformed the micro-cavity surface (**Supplementary Fig. 5**). No deformation was observed for T cells on uncoated and on type 1 collagen-coated micro-cavities, indicating that the surface interactions are protein specific. By analysing a large number of indentation events, we found that the strength of adhesion increased with the surface concentration of ICAM-1 as more LFA-1:ICAM-1 complexes can be formed at higher ICAM-1 surface concentration.

### Horizontal traction stresses in mesenchymal cell migration

Next, we investigated cells that are known to apply chiefly horizontal traction forces to their substrate when in a two-dimensional environment. Although ERISM performs direct measurements of vertical deformations of the micro-cavity substrate, it is also highly sensitive to horizontal forces. Applying a defined lateral force to the surface of an ERISM micro-cavity using AFM led to local twisting of the top gold surface. The amount of twisting was directly proportional to the applied force (150 pN per nm of twist; *R*^2^ > 0.99; *n* = 5). The origin of this twisting is somewhat related to but different from the non-linear surface wrinkling effect that early studies on cell generated traction forces have used^20,21^: The interferometric detection principle of ERISM provides sufficient sensitivity to detect small and localized, linear surface twisting; taking into account the 2 nm displacement resolution of ERISM, the minimum lateral force that causes a clearly resolvable signal is < 300 pN (**Supplementary Fig. 6**).

**Figure 4a** shows a phase contrast image of a 3T3 fibroblast attached to the surface of a micro-cavity and the corresponding ERISM map. The cell was fixed during the measurement and then immunolabelled for actin and vinculin (**Fig. 4b**). **Figure 4c** shows the Fourier filtered ERISM map of the cell in which vinculin-rich regions are highlighted. At the periphery of the cell, there were multiple clearly distinguishable push/pull features (i.e. twisting; marked by pairs of blue/red arrows in the magnified inset to **Fig. 4c**). This twisting is consistent with earlier observations of torque being applied by focal adhesions^26^. Areas of high vinculin expression were located at the centre of these features, confirming that the observed features are associated with focal adhesions that transfer traction forces horizontally to the substrate. Focal adhesion complexes had sizes of (6.1±3.4) μm^2^ and transmitted horizontal forces and stresses of (3.1±1.0) nN and (602±323) Pa, respectively, to the substrate (*n* = 19). These values are not significantly different from reference measurements we performed using conventional TFM on stiffness-matched polyacrylamide gels (horizontal stress at focal adhesion, (386±131) Pa; *n* = 6; *p* = 0.11, Mann-Whitney *U*- test). They are also consistent with earlier micro-pillar array measurements of the force transmitted by focal adhesions of Bovine pulmonary artery smooth muscle cells (≈20 nN horizontal force applied by 4 μm^2^-sized adhesion complexes on an array with fivefold larger effective stiffness)^29^ and with TFM measurements of mouse embryonic fibroblasts (≈500 Pa horizontal stress on a hydrogel substrate with a shear modulus of ≈2.4 kPa)^26^.

**Figure 4.**
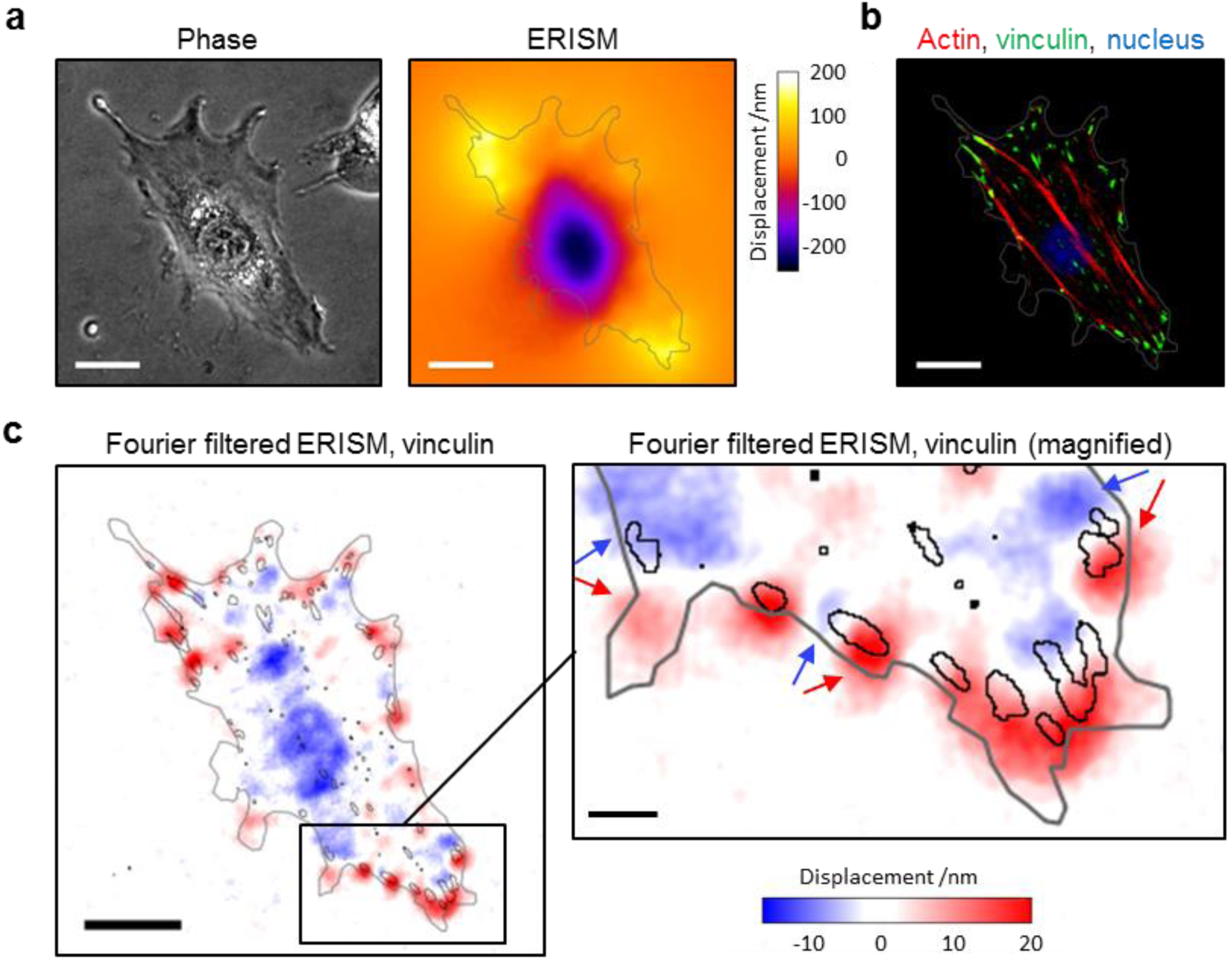
Analysis of the horizontal stress applied by adherent 3T3 fibroblasts using ERISM. (a) Phase contrast image and false colour map of the micro-cavity deformation generated by a 3T3. The cell boundary is shown in grey. (b) Epi-fluorescence image of same cell following fixation and staining for actin, vinculin and nuclear DNA. (c) Fourier filtered ERISM data from (a). Cell outline in grey. Black lines indicate vinculin-rich areas, overlay of data from (b). All scale bars 25 μm; except for inset to (c) 5 μm.

### Long-term imaging of mechanical activity: cell cycle, migration from tissue explants and *in situ* differentiation

We then applied ERISM to monitor cell behaviour with high temporal resolution and over extended periods of time. Time-lapse data of slowly migrating 3T3 fibroblasts with a frame taken every 2 s revealed fast force oscillations (oscillation period, 11.3 s; **Supplementary Video 5** and **Supplementary Fig. 7**), which is consistent with earlier observations^5^. **Supplementary Video 6**, on the other hand, shows a 5.5 day time-lapse video of migrating 3T3 fibroblasts with a frame taken every 5 minutes and a total of over 1,600 frames. This measurement shows five cycles of cell division with no signs of phototoxicity or micro-cavity degradation, thus enabling the analysis of the mechanical activity of cells during and between mitotic events. During mitosis, cells rounded up which was associated with a complete loss of applied stress (**Fig. 5a**), consistent with earlier observations^21^. Following cell division, cells quickly reattached and began to apply forces to the substrate. Taking the total volume by which each cell indents into the substrate as a proxy for the applied force, we observed pronounced force fluctuations (period of oscillation, (21.5±5.0) min; **Fig. 5b**). We found the amplitude of these fluctuations to increase quickly after each cell division, before going through a maximum followed by a slow decay in force as the cells approached mitosis again (**Fig. 5c**).

**Figure 5.**
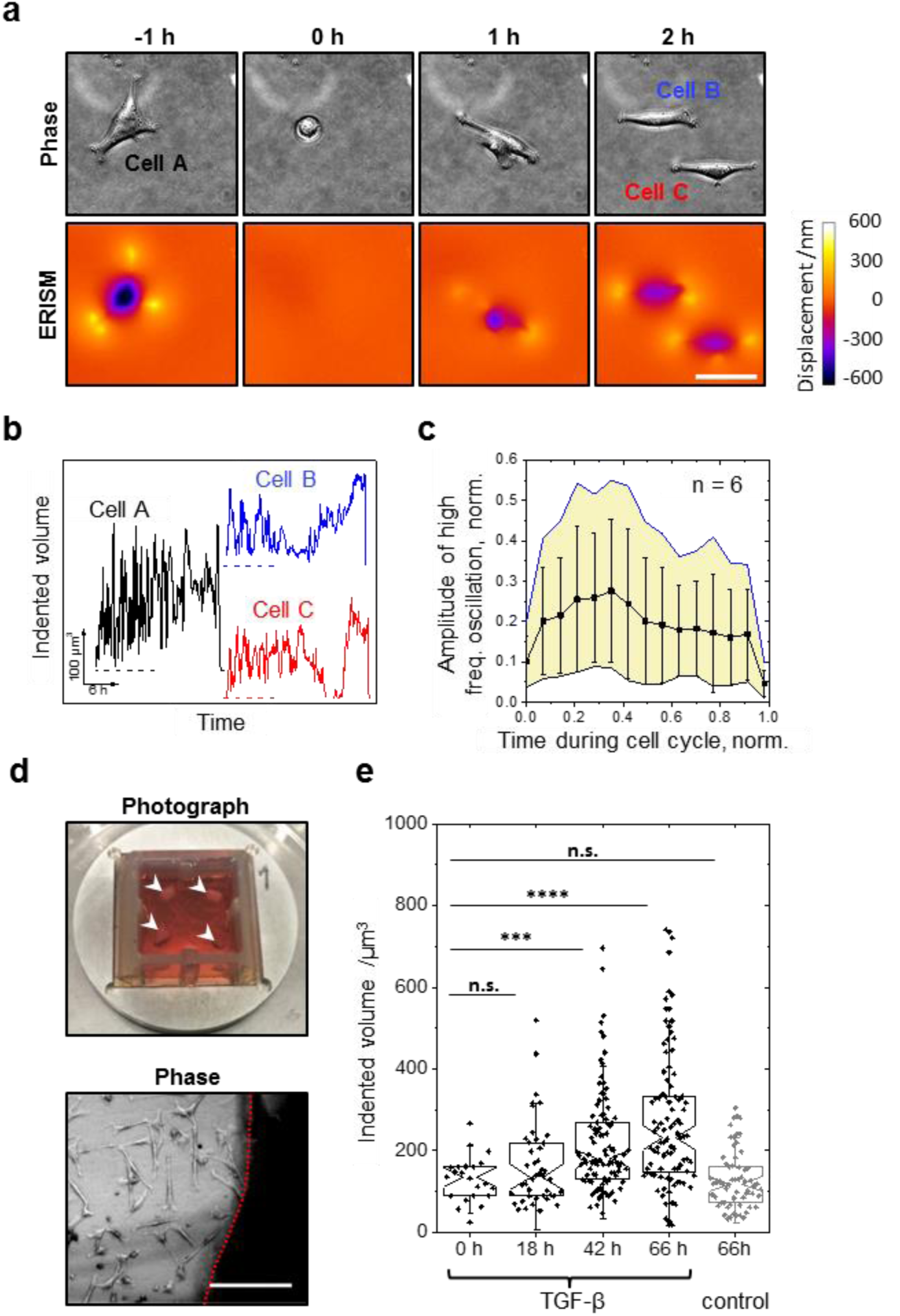
ERISM for long-term time lapse studies of the mechanical activity of cells. (a) Representative phase contrast and ERISM images from a 1,600 frame time-lapse series which recorded the mechanical activity of 3T3 fibroblasts over 5.5 days. Shown is one of several cell-division events with mother cell (Cell A) and two daughter cells (Cell B and C). (b) Temporal evolution of the mechanical activity of Cells A-C over the course of their cell cycles. Traces show a high-frequency oscillatory behaviour that changes in amplitude over time. (Traces vertically displaced for clarity; dashed lines indicate zero levels for each trace.) (c) Analysis of the oscillatory mechanical activity of *n* = 6 fibroblasts during their cell cycle. Shown are the mean (black squares), standard deviation (whiskers) and minimum and maximum amplitude (yellow area) of the oscillation amplitude. (d) Photograph of four pieces of mouse skin (marked by arrow heads) incubated directly on a micro-cavity substrate and bright-field microscopy image of primary fibroblasts migrating out of the skin (shadowed area separated by red dashed line) on day four of incubation. (e) Investigation of TGF-β-induced differentiation of primary mouse fibroblasts into myofibroblasts. Tukey boxplot of mechanical activity before (0 h) and during (18 h, 42 h and 66 h) incubation with 10 ng mL^-1^ TGF-β and with media alone (control group). Each dot represents one cell. As data in groups was not normally distributed, groups were compared using the Mann-Whitney U-test (\*\*\**p* < 0.001; *\*\*\*\*p* < 0.0001). Scale bars, 50 μm in (a) and 200 μm in (d).

Next, we applied ERISM to primary mouse fibroblasts. The robustness of the ERISM micro-cavity allowed for tissue explants to be incubated directly on the cavity for several days until cells had migrated out and onto the micro-cavity substrate (**Fig. 5d**). As zero-force reference images are not required for data analysis, we were able to record the mechanical activity of fibroblasts in close proximity to the tissue explant (**Supplementary Videos 7 and 8** and Supplementary **Fig. 8**). We then stimulated the fibroblasts with TGF-β to initiate their differentiation into myofibroblasts^10^. Myofibroblasts express large amounts of a-smooth muscle actin (a-SMA) and play a critical role in wound healing; they accumulate in the wound’s granulation tissue and contribute to wound closure by applying large contractile forces^10^. **Figure 5e** shows the analysis of the cellular forces exerted during the differentiation process (see **Supplementary Fig. 8** for ERISM map and a-SMA immunostaining data for independent confirmation of differentiation). Over the 66 h time course of the experiment, the mean force of cells stimulated with TGF-β increased by about twofold, whereas no change in force was observed for the control group. ERISM allows sampling large numbers of cells from the same population at different time points over the course of an experiment (in total, more than 300 cells were measured and analysed in this example), which is beneficial for statistical analysis.

## Discussion

ERISM is a fast, robust and sensitive method for recording mechanical cell-substrate interactions that is based on analysing local changes in the resonance wavelengths of an elastic, semi-transparent optical micro-cavity. It can be used to measure forces <1 pN that cells apply perpendicular to the substrate surface with a spatial resolution < 2 μm, and it also readily detects lateral forces with < 300 pN sensitivity. We validated ERISM by AFM, using indentation and horizontal pulling measurements, and applied it to a wide range of biological systems. ERISM can be easily combined with other microscopy modalities, such as bright-field, phase contrast or fluorescence microscopy. The micro-cavity substrates used for ERISM show excellent long-term stability under mechanical stress and standard cell culture conditions, which renders ERISM particularly well suited for measuring cellular forces over prolonged periods of time. The extraction of cells from primary tissue directly on a micro-cavity facilitated the investigation of primary cells without transferring them between substrates. The fact that no zero-force reference image is required to analyse ERISM data eliminates the need to detach non-migrating cells after a measurement, which makes them available for subsequent studies (e.g. by immunostaining) and drastically simplifies investigation of large numbers of cells on a common substrate and at multiple time-points. To illustrate this capability, we measured forces of more than 300 cells during the differentiation of fibroblasts into myofibroblasts and found a twofold increase in cellular force, in line with earlier TFM measurements on a sample of much smaller size^44^.

Methods to measure cellular forces that require high light intensities can be problematic in terms of cellular photo-damage, in particular if repeated measurements are required during long-term investigations. The intensity needed to acquire the reflectance images in ERISM is in the 100 μW cm^-2^ range – much lower than the intensities required for fluorescence-based methods such as TFM. Furthermore, because the interference effect occurs within the substrate and does not require an external reference mirror, ERISM time-lapse data is free from drift-artefacts. Together, these features enable reliable long-term time-lapse measurements, as demonstrated by a 5.5 day continuous time-lapse consisting of over 1,600 frames.

The extreme stress sensitivity, big field of view, large dynamic range, and the capability to perform continuous long-term measurements provide unique possibilities for observing podosome protrusions during cell migration – the process during which podosome activity is believed to be most crucial. In addition, ERISM revealed that the migration of *Dictyostelium discoideum* under confined conditions differs fundamentally from the situation on flat substrates, for which earlier TFM studies found large in-plane stresses exerted by stationary traction adhesions^27,39,45,46^. Like many other cell types, *Dictyostelium discoideum* is capable of adapting its mode of locomotion to its environment^18,40,47-49^. Our data suggest that in a spatially confined environment they project themselves forward in horizontal direction by applying weak vertical forces, a mechanism previously described as “chimneying” ^18,38,41^. A recent investigation on non-adherent Walker 256 carcinosarcoma cells found that friction stresses play a pivotal role for migration through micro-channels^17^. While these friction stresses were too small to be resolved with TFM, they were estimated to be comparable to the stresses we measured with ERISM for *Dictyostelium discoideum*, indicating that ERISM may be well suited to study cell migration in micro-channels and other confined environments.

In summary, ERISM provides a direct and simple measurement of cellular forces. Its field of view and lateral resolution can be readily adjusted by changing the microscope objective. The shape of the FEM-reconstructed stress maps largely follows the displacement profiles so that approximate stress information can be obtained by simply counting interference fringes in a single monochromatic reflectance image. Finally, the optics needed for ERISM can be easily integrated into a standard inverted cell culture microscope. We anticipate that the ability of ERISM to quickly assess normal versus abnormal mechanical behaviour of cells, together with the option to investigate multiple cells within a large field of view online, will render ERISM a very powerful tool for diagnostic applications.

## Acknowledgments

The authors thank K. Venkatesan Iyer (Max Planck Institute for Cell Biology and Genetics, Germany) and Paul Reynolds (University of St Andrews, U.K.) for fruitful discussion, Rajesh Shahapure for TFM reference measurements, Anna L. Sobiech for illustrations and the DictyoBase for provision of AX3 strain Dictyostelium discoideum. This work was supported by the Human Frontiers Science Program (RGY0074/2013), the Scottish Funding Council (via SUPA), the EPSRC DTP (EP/L505079/1), the RS MacDonald Charitable Trust and the MRC project (G1100116).

## Author Contributions

N.M.K., P.L. and M.C.G. developed ERISM. N.M.K. fabricated and characterized the micro-cavity substrates and conducted measurements. P.L. developed the data analysis, stress map calculation and graphical data presentation. A.S. contributed to protein coating and staining and supported cell culture. J.A.K. designed and prepared the primary mouse fibroblast experiment. J.G.B. prepared and assisted the T cell experiment. G.S. and K.F. performed part of the initial elastomer characterization through rheometry and AFM. S.J.P. proposed and prepared the macrophage experiment. M.C.G. supervised the project. N.M.K. and M.C.G wrote the manuscript with input from all authors.

## Online Methods

### Device fabrication

The bottom mirror of the micro-cavity substrate is formed by a three-layer stack comprising a 0.5 nm-thick chrome adhesion layer, a 10 nm-thick Au mirror layer and a 50 nm-thick SiO_2_ layer. Layers were deposited by electron-beam evaporation onto cleaned 2.4 cm x 2.4 cm cover slips (500-600 μm thickness). The two components of the siloxane-based elastomer (GEL8100, NuSil) were thoroughly mixed in a 1:1 volume ratio and degassed under vacuum for 1 min. The mixture was spin coated onto the bottom cavity mirror to yield 5 μm-thick films (1 min at 5000 rpm, measurement of *Dictyostelium discoideum)* or 8.0 μm-thick films (1 min at 3000 rpm, all other cells) and crosslinked on a hotplate at 110 °C for 1 h. The surface of the elastomer layer was exposed to mild oxygen plasma treatment (power, 15 W; duration, 30 s) to increase its surface energy (**Supplementary Fig. 1**). Finally, 15 nm of Au are thermally evaporated to complete the optical cavity.

### Cell culturing and cell-mechanical investigations

For cell culturing a silicone chamber (Ibidi) with inner dimensions of 0.75 x 0.75 cm^2^ or 1.60 x 1.60 cm^2^ (for tissue explant experiment) was mounted onto the micro-cavity. The substrate was incubated with a type I collagen suspension (Collagen A, Biochrome) at pH 3-3.5 over night and then washed with cell culture medium. For T cells, micro-cavities were incubated with 260 μL or 350 μL of 3 μg mL^-1^ ICAM-1 (R&D Systems) in PBS overnight at 4 °C, to achieve the different ICAM-1 surface concentrations.

For isolation of human macrophages, peripheral blood was obtained from a normal healthy donor after ethical review (School of Medicine, University of St Andrews) and informed consent. Peripheral blood mononuclear cells were purified over a Ficoll gradient and allowed to adhere to plastic culture dishes at 37°C for 60 mins in RPMI 1640 (Invitrogen) supplemented with 5% FCS (Invitrogen). After extensive washing to remove lymphocytes, the adherent monocytes were cultured in 50 ng ml^-1^ GM-CSF (Immunotools) to generate macrophages. Flow cytometry was performed by blocking the cells in 10% human serum followed by incubation with an irrelevant antibody or anti-CD14-PE (eBioscience, UK) (see **Supplementary Fig. 9**). The cells were 90% CD14^+^. Cells were transferred to a micro-cavity substrate in serum free RPMI which promoted surface adhesion.

T cell suspensions were prepared from lymph nodes of Rag1-/-OT-I TCR transgenic mice. Biotinylated Abs were used for purification of naive CD8 T cells by negative selection: anti-CD4-Pe (eBiosciences), anti-I-A/I-E-Pe (Biolegend) followed by a 15-min incubation with anti-Pe MACS beads (Miltenyi Biotech). Cells were added to MACS LS columns (Miltenyi Biotech) and unstained CD8 T cells collected. CD8 T cells were maintained in IMDM media (Invitrogen) supplemented with 5% FCS, L-glutamine, antibiotics and 50 μM 2- mercaptoethanol. CD8 T cells were activated with 5 μg mL^-1^ soluble anti-CD3 Ab (clone 2C11, R&D Systems) immediately after they were placed on the micro-cavity substrate. Cells were incubated for 10 min prior to ERISM.

AX3 strain *Dictyostelium discoideum* amoebae (Dictybase) were cultured in HL-5 medium (Formedium). A spatially confined environment above the micro-cavity substrate was generated by dispensing 10 μm-diameter glass spheres on a 5 μm-thick micro-cavity (without collagen coating). AX3 cells were dispersed on the chip in 1 x PBS buffer solution and the droplet was gently spread over the substrate by placing a 500 μm-thick glass coverslip (without collagen coating) onto the glass spheres. A 5 μm-void was formed between the top surface of the micro-cavity and the bottom surface of the glass coverslip.

3T3 fibroblasts (Sigma Aldrich) were seeded on the substrate in DMEM medium (Gibco, with GlutaMAX) with 10vol% fetal calf serum (FCS, Biochrom) and 1vol% penicillin-streptomycin solution (PS, Gibco) and cultivated for 24 hours before performing ERISM.

Primary mouse fibroblasts were obtained by culturing skin explants from a Balb/c mouse on the micro-cavity substrate. The Balb/c mouse was bred in-house; animal housing conditions were approved by the University of Edinburgh and in accordance with the Animals (Scientific Procedure) Act 1986. Mouse skin was sterilised using antiseptic solution (Videne, Ecolab) and washed in PBS. Skin pieces (2 x 2 mm^2^) were adhered to the surface of the micro-cavity with the epidermis side pointing upwards and incubated in DMEM medium containing 10vol% FCS and 1vol% PS. Primary fibroblasts were differentiated into a myofibroblastic phenotype using 10 ng mL^-1^ recombinant mouse TGF-β1 (RnDSystems). For α-SMA staining, cells were fixed directly on the micro-cavity substrate with 4% paraformaldehyde (Alfa Aesar) in PBS at room temperature for 15 min, washed with 0.05% Tween 20 in PBS (Alfa Aesar), permeabilized with 0.1% Triton X-100 (Alfa Aesar) for 3 min and blocked for 30 min with 1% BSA (Carl Roth) in PBS. Cells were stained using a mouse anti-rabbit-α-SMA monoclonal antibody (Sigma-Aldrich, 1:500, for 1h at room temperature), followed by the secondary goat anti-mouse FITC-conjugated antibody (Sigma Aldrich, 1:32 for 30 min).

For time lapse ERISM measurements, the micro-cavity substrate was transferred into an on-stage microscope incubator (OKOLAB) providing standard cell culture conditions (37 °C, 5vol% CO_2_, 100% humidity) over extended durations. *Dictyostelium discoideum* cells were measured at room temperature.

For staining of macrophages and 3T3, cells were fixed directly on the micro-cavity substrate with 4% paraformaldehyde (Alfa Aesar) in PBS at room temperature for 20 min, washed with 0.05% Tween 20 in PBS (Alfa Aesar), permeabilized with 0.1% Triton X-100 (Alfa Aesar) for 3 min and blocked for 30 min with 1% BSA (Carl Roth) in PBS. Cells were stained for vinculin using a mouse anti-vinculin monoclonal antibody (Merck Millipore, 1:250 in BSA solution, 1h at room temperature), followed by the secondary goat anti-mouse FITC-conjugated antibody (Sigma Aldrich, 1:32 in BSA solution, 30 min). Actin was stained in parallel using TRITC-conjugated Phalloidin (Merck Millipore, 1:500 in BSA solution, 1h at room temperature). Cell nuclei were stained with DAPI (Merck Millipore, 1:1000 in BSA, 3 min at room temperature).

### ERISM measurement

The local resonance wavelengths of the micro-cavity were determined by imaging its reflectance at 201 different wavelengths between 550 nm and 750 nm in epi-illumination configuration. Monochromatic probe light (bandwidth, 4-5 nm full width at half maximum) was provided by a halogen lamp and a 1/8 m monochromator with a 1200 mm^-1^ grating (blaze wavelength, 650 nm) and 0.6 mm-wide slits (CM110, Spectral Products). The probe light was coupled to an inverted fluorescence microscope (Nikon Ti-S with stratum structure) and focussed onto the rear focal plane of a 10x, 20x, or 40x objective by custom-built Köhler-like illumination optics to achieve high contrast and low background during the measurement. The interference patterns reflected by the micro-cavity were imaged through the same objective and recorded with a sCMOS detector (Zyla 5.5, Andor Technology). Wavelength scanning and imaging were controlled by custom-written software (LabView, NI). Each full wavelength scan (550-750 nm) took 8-20 s depending on exposure settings. The optical readout time can be reduced (down to 2 s in our present system configuration) by performing the full wavelength scan only once to find the resonance order of the interference fringes. Subsequent measurements then track the spectral position of only one interference minimum, which reduces the required scan range to 30-50 nm.

A second camera (mounted to the auxiliary microscope port) was used to acquire bright-field, phase contrast or fluorescence images. For time-lapse measurements, both cameras were controlled by the custom-written software.

### Data analysis

The local resonance wavelengths were extracted at every position across the field of view by analysing the wavelength-dependence of the reflectance for each of the 1.3 million pixels in the image stack. For each pixel the reflectance was analysed as a function of wavelength and the minima were identified. The thickness for each pixel was determined by matching the measured local resonance wavelengths against a pre-compiled database containing the resonance wavelengths for all possible cavity thicknesses. The pre-compiled database was generated by transfer matrix method calculations^1^ and the wavelength matching was performed with a fast least-absolute-deviation fitting algorithm. When allowing for a maximum deviation of 1 nm per resonance wavelength, the algorithm typically yielded less than 0.01% of non-fitted pixels across the entire field of view. In some cases, the absolute deviation between measurement and simulation was almost equally small for thicknesses associated with different interference orders. We therefore applied a thickness continuity criterion (maximum thickness change between pixels <50 nm) to eliminate unphysical jumps in thickness. Finally, a background plane was subtracted from the measured thickness map to obtain the relative micro-cavity deformation. The entire data analysis is fully automated and is based on custom-written software (Cython / Python, Python Software Foundation). We have not rigorously optimized our algorithm for speed and with the current implementation each deformation map requires about 5 s of computation time at full resolution (1280 x 1024 pixels) on a standard desktop PC. For FF-ERISM, an FFT bandpass filter was applied to the raw displacement maps using the ImageJ software.

### Atomic Force Microscopy (AFM) indentation measurements

AFM (FlexAFM, Nanosurf) indentation measurements were carried out in concentrated buffer solution (CHAPS, 3-[(3-cholamidopropyl)dimethyl ammonio]-1-propane sulphonate, Roth) to minimize electrostatic interaction between AFM probe and sample. Spherical glass beads with a diameter of 10 μm or 17 μm were glued to cantilevers with nominal force constants of 0.01, 0.1 and 0.3 N m^-1^. (Actual force constants were individually measured by the thermal-tuning-method before attaching the bead.) The cantilever deflection was calibrated by pushing the beaded cantilevers against a rigid glass substrate using a known z-travel distance (typically 0.5 μm). Force-distance curves were obtained for indentations into the elastomer layers or micro-cavity and analysed by fitting with a corrected Hertz model^2^ to obtain the apparent stiffness of the sample. The Poisson’s ratio of the elastomer was assumed to be 0.49, a value previously reported for the structurally similar elastomer PDMS^3^.

The corrected Hertz model assumes a homogeneous, linear elastic material and no friction between indenter and sample surface. These assumptions are not fulfilled for the layered architecture of our micro-cavities and the slight deviation between the measured force-distance curve and the Hertz fit for the micro-cavity in **Fig. 1h** of the main manuscript is attributed to this effect. We therefore refer to the Young’s moduli derived from corrected Hertz model fits as the apparent stiffness, *E\**.

Lateral AFM measurements were performed with a tipped cantilever and without the CHAPS buffer so that electrostatic attraction fixed the cantilever on the surface of a microcavity. No slipping of the tip was observed. The lateral bending stiffness of the cantilever used was 62 N/m (taking into account torsion and lateral bending). Horizontal force was applied by moving the in-plane scan-head piezo of the AFM.

### Calculation of stress map and exerted cellular force from displacement map

Local vertical stresses were derived from the displacement maps using FEM (COMSOL Multiphysics 5.0). The oxide and gold layer on top of the elastomer layer slightly broaden stresses applied to the surface of the micro-cavity but do not reduce the total force applied. We therefore modelled the micro-cavity as an isotropic linear elastic material with thickness of 8.5 μm, a Young’s modulus of 0.30 kPa (see **Supplementary Fig. 10**) and a Poisson’s ratio of 0.49. By applying the measured displacement to the top surface of an FEM model of the structure, the Cauchy stress tensor at the interface between elastomer and top layer was obtained and its vertical component was then used as a measure of the local stress applied by cells. Total forces were obtained by integrating over the displaced area in the corresponding quantitative stress maps using measurement routines in the ImageJ software.

